# Cellular crowd control: overriding endogenous cell coordination makes cell migration more susceptible to external programming

**DOI:** 10.1101/2021.01.23.427700

**Authors:** Gawoon Shim, Danelle Devenport, Daniel J. Cohen

## Abstract

As collective cell migration is essential in biological processes spanning development, healing, and cancer progression, methods to externally program cell migration are of great value. However, problems can arise if the external commands compete with strong, pre-existing collective behaviors in the tissue or system. We investigate this problem by applying a potent external migratory cue—electrical stimulation and electrotaxis—to primary mouse skin monolayers where we can tune cell-cell adhesion strength to modulate endogenous collectivity. Monolayers with high cell-cell adhesion showed strong natural coordination and resisted electrotactic control, with this conflict actively damaging the leading edge of the tissue. However, reducing pre-existing coordination in the tissue by specifically inhibiting E-cadherin-dependent cell-cell adhesion, either by disrupting the formation of cell-cell junctions with E-cadherin specific antibodies or rapidly dismantling E-cadherin junctions with calcium chelators, significantly improved controllability. Finally, we applied this paradigm of weakening existing coordination to improve control to demonstrate accelerated wound closure *in vitro*. These results are in keeping with those from diverse, non-cellular systems, and confirm that endogenous collectivity should be considered as a key, quantitative design variable when optimizing external control of collective migration.

## Introduction

Collective cell migration enables intricate, coordinated processes that are essential to multicellular life, spanning embryonic development, self-healing upon injury, and cancer invasion modes^1^. Control of collective cell migration, therefore, would be a powerful tool for biology and bioengineering as such control would enable fundamentally new ways of regulating these key processes, such as enabling accelerated wound healing. Efficient and precise control over cell motility is becoming increasingly feasible with modern biotechnologies. Tunable chemical gradient generators can redirect chemotaxing cells^2,3^, optogenetics can allow dynamic control of cell contractility^4^, micropatterned scaffolds can constrain and direct collective growth^5^, and recent work in bioelectric interfaces has even demonstrated truly programmable control over directed cell migration in 2D^6,7^. However, despite advances in sophisticated tools, applying them to complex, cellular collectives raises a fundamental problem: what happens when we command a tissue to perform a collective behavior that competes with its natural collective behaviors?

Paradoxically, those endogenous collective cell behaviors already present in tissues are both a boon and bane for attempts to control and program cell behavior. On the one hand, endogenous collective cell migration means the cells already have established mechanisms for coordinated, directional migration that external cues and control can leverage. For instance, cadherin mediated cell-cell adhesions in tissues mechanically couple cells together and allow for long-range force transmission and coordinated motion. This coupling allows tissues to migrate collectively and directionally over large distances and maintain cohesion and organization far better than individual cells might^8,9^. On the other hand, imposing a new behavior over an existing collective behavior may generate conflicts. Tight cell coupling can create a ‘jammed state’ or homeostatic tissue where cells are so strongly attached and confined that they physically lack the fluidity to migrate as a group^10,11^. Strong coordination established via physical coupling can hinder cells from responding to signals for migration, as shown by the need for zebrafish and other embryos to weaken cell-cell junctions prior to gastrulation to ensure cells collectively migrate to necessary locations^12–14^. Hence, how ‘susceptible’ a collective system may be to external control likely depends on a tug-of-war between the resilience and strength of the natural collective processes and the potency of the applied stimulus.

Here, we specifically investigate the relationship and interplay between an applied, external command attempting to direct collective cell migration, and the strength of the underlying collective behaviors already present in the tissue. We address two key questions: 1) how much does the strength of an endogenous collective migration behavior in a tissue limit our ability to control its collective cell migration, and 2) how can we circumvent such limitations? To investigate these questions, we needed both a programmable perturbation capable of controlling collective migration, and a physiologically relevant model system allowing for tunable ‘collectivity’. As a perturbation, we used the SCHEEPDOG bioreactor^6^ to harness a bioelectric phenomenon called ‘electrotaxis’—directed cell migration in DC electric fields—which can broadly induce large-scale directional migration *in vitro* in over 20 cell types and is implicated in a number of developmental processes as a navigational cue guiding cell migration *in vivo*^15–17^. Briefly, electrotaxis arises when endogenous, ionic fields form during healing or development (∼1 V/cm) and apply gentle electrophoretic or electrokinetic forces to charged receptors in cell membranes, causing them to aggregate towards one side of a cell and produce a front-rear polarity cue^18,19^. Electrotaxis is perhaps the only cue that can guide large-scale migration in a broad range of cell and tissue types without any modifications, and this generality and prior demonstrations of collective electrotaxis^6,8,20^ made it a strong candidate.

To complement electrotaxis, we chose primary mouse skin for our model system as skin injuries were where the endogenous electrochemical fields that cause electrotaxis were first discovered (center of a wound is negative relative to the periphery), and we and others have shown layers of keratinocytes to exhibit strong electrotaxis^6,21–23^. Critically, primary mouse keratinocytes have tunable ‘collectivity’ in culture. Specifically, cell-cell adhesion strength in this system, mediated by cadherin proteins, can be easily tuned by varying calcium levels in the media—with low calcium media thought to mimic conditions in the basal layers of the epidermis with weak adhesions, and high calcium media akin to conditions in the uppermost layers of skin with strong adhesions^24–27^.

Together, these experimental approaches allowed us to precisely explore how the ability to externally ‘steer’ collective migration in a living tissue using a powerful bioelectric cue depends on the native collectivity of the underlying tissue. First, we quantify and validate that we can tune collective strength in cultured skin layers, and link collectivity to E-cadherin and collective migration phenotypes. Next, we demonstrate how applying the same electrical stimulation conditions to tissues with differing native collectivity results in radically different outputs with weakly collective tissues precisely responding to our attempts to control their motion, while strongly collective tissues exhibited detrimental supracellular responses resulting in tissue collapse. We then prove that E-cadherin is responsible for these differences, ruling out any effects of calcium signaling per se. Finally, we leverage these findings to develop a new approach that allows us to effectively control mature, strongly collective tissues, which we utilize to demonstrate that we can accelerate wound repair *in vitro*.

## Results

### Establishing baseline collective migration of primary keratinocyte layers

To determine how natural collective cell behaviors compete with externally imposed control of collective behavior, we first need to establish baseline data of endogenous collective behavior in the absence of guidance cues. We used layers of mouse primary keratinocytes as a model system as their endogenous collective behavior is well-characterized^22,28^, they have a strong electrotactic response^6^, and their cell-cell adhesion levels can be easily tuned via calcium levels in the culture media^27,29^. Previous work has indicated that cell-cell adhesions via calcium-dependent proteins, E-cadherin adhesion being one of the best-studied, are essential in interconnecting individual cells and maintaining coordination within the monolayers by coupling mechanical information via the cadherin-catenin-actin complex^30–33^. Hence, we hypothesized that modulation of cell-cell adhesion levels via calcium control would allow us to tune the relative strength of collective couplings and collective migration in primary keratinocyte layers, giving us a precise and reproducible system to explore questions of collective control.

To establish quantitative standards for collective strength in our keratinocyte model, we engineered arrays of identical 2 x 2 mm keratinocyte tissues using tissue stenciling methods^6,34^. Tissue arrays were then cultured for 14 h in high (1.0 mM), medium (0.3 mM), or low (0.05 mM) calcium conditions to allow junction formation (Fig. 1A). These calcium levels are standard conditions that span the physiological range based on phenotypes and marker expressions^27,29,35,36^. As E-cadherin is a major calcium-dependent adhesion protein, we used immunostaining to quantify and confirm the direct relationship between calcium level and E-cadherin recruitment to cell-cell junctions (Figs. 1A, 1B, S1). We generated collective migration data for each tissue by processing phase-contrast timelapse movies captured using automated imaging (Methods) with Particle Image Velocimetry (PIV) to generate velocity vector fields at each time point. The vector fields were then analyzed to visualize and quantify the strength of coordinated motion within a given tissue over time (Fig. S2)^6,21,34^. First, we calculated the directionality of cellular movements to visualize domains of coordinated migration within tissues. Directionality (Eqn. 1) is defined as the average of the cosine of the angle between each PIV velocity vector and the horizontal x-axis, while N is equal to the total number of velocity vectors in the frame. As the electric field command is also in the horizontal direction, the directionality also indicates how well aligned the cellular migration is with the field direction under stimulation. Directionality can vary between -1 (cell motion to the ‘left’; perfectly anti-parallel with field) and 1 (cell motion to the right; perfectly parallel with field). Additionally, we quantified the collectivity by calculating the overall coordination within a tissue using the polarization order parameter (Eqn. 2) from collective theory, where *ν*_*i*_ indicates the *i*th velocity vector^37^. A coordination value of 1 indicates perfect coordination and anistropy across the whole tissue, while 0 indicates wholly isotropic motion.

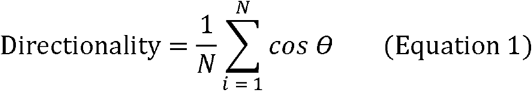

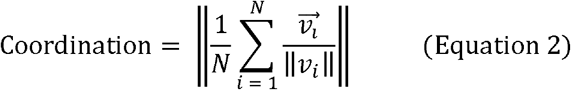

Our data (Figs. 1C, 1D) clearly demonstrate that increasing calcium levels increases collectivity within the tissue. Both the general size of coordinated domains, represented by large zones of either red or blue in Fig. 1C, and the coordination parameter varied directly with calcium levels (Fig. 1D). The Velocity correlation function for nearest neighbors also show higher correlation with increased calcium levels (Fig. S8. However, we also noted that increased coordination came at the cost of reduced average cell migration speed (Fig. 1E, Movie S1), suggesting that strong cell-cell adhesion impeded cellular motion, a common tradeoff in collective motion^38^. Notably, there is a clear shift in cell and tissue morphology across the different calcium levels, with high calcium tissues visually exhibiting supracellular fluctuations and low calcium tissues behaving far more like a dense collection of individualistic agents. Together with our data indicating that E-cadherin levels also vary directly with calcium, and prior studies indicating a strong correlation between cadherin levels and coordination, these data validated our ability to tune endogenous collective strength in keratinocyte layers, and to quantify and profile the natural collective motion of unstimulated tissues. With baselines established, we next investigated how collective strength regulates electrotactic susceptibility.

**Figure 1.**
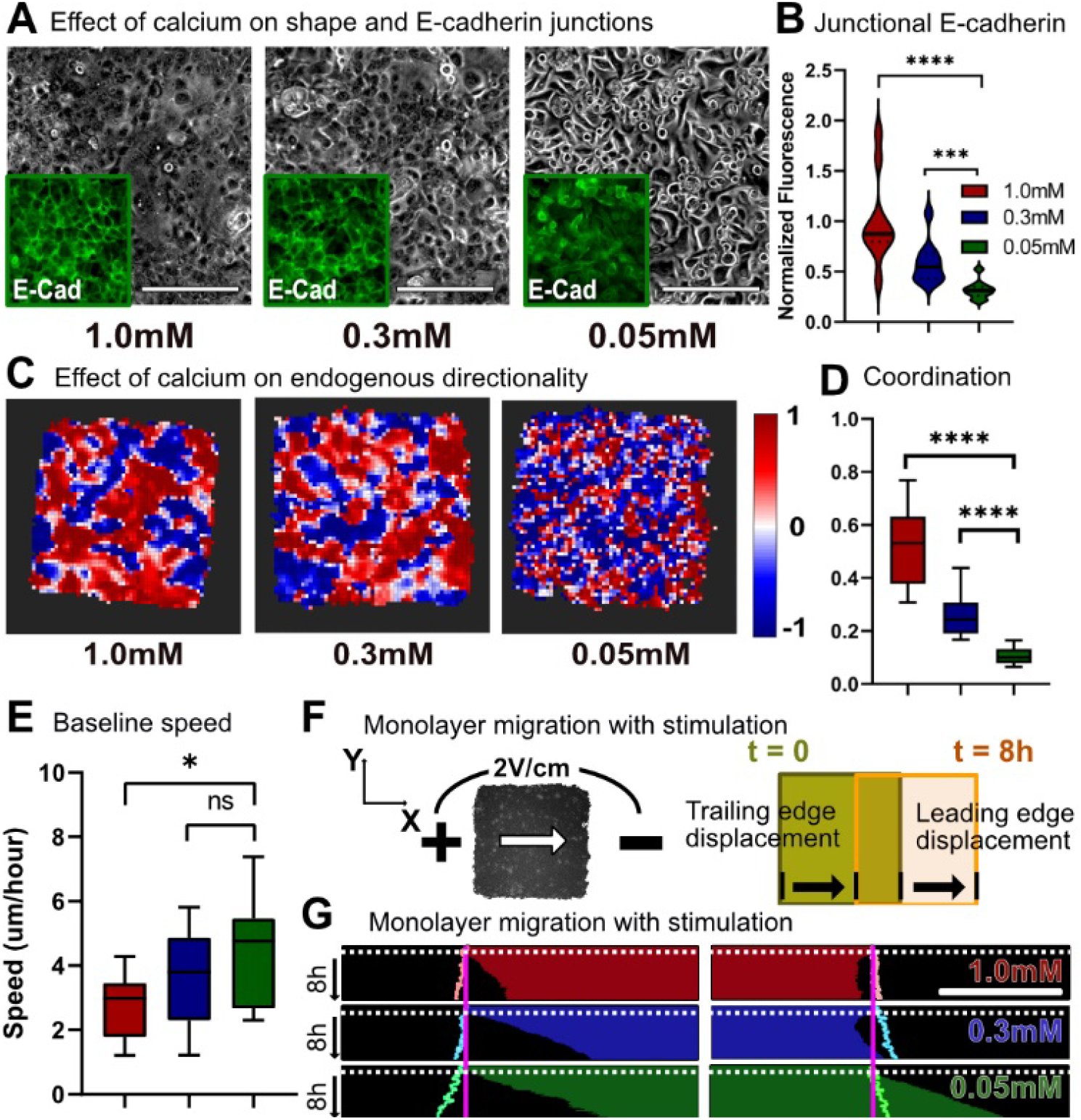
Baseline collective behavior of keratinocyte monolayers; endogenous coordination increases with calcium-dependent cell-cell adhesion. A) Phase and E-cadherin imaging for primary mouse keratinocyte monolayers cultured in high (1.0mM), medium (0.3mM), and low (0.05mM) calcium media for 14h. Grey: phase image; green inset: immunofluorescence image of E-cadherin. Scale bar = 200um. B) Distribution plot for the normalized junctional E-cadherin immunofluorescence signal for high, medium, and low calcium monolayers. C) Horizontal directionality heatmap for monolayers of varying calcium. D) Distribution plot for coordination values for monolayers of varying calcium. Legends identical to B). E) Baseline migration speed for monolayers of varying calcium. F) Schematic for keratinocyte monolayer migration towards the cathode; leading and trailing edge displacement. G) Leading and trailing edge displacement kymographs for monolayers of varying calcium throughout 1h control (no stimulation) and 8h stimulation. Electrical stimulation starts at white dotted line. Pastel outlines indicate the edge displacement of unstimulated monolayers at same calcium level throughout 9h. Scale bar = 500um. P values are calculated using unpaired nonparametric Mann-Whitney test with n = 15 for each condition. ** corresponds to p < 0.001, and **** to p < 0.0001.

### Strong collectivity makes it more difficult to program collective cell migration

Having related low calcium levels to weak collectivity and low junctional E-cadherin, and high calcium levels to strong collectivity and high junctional E-cadherin, we next attempted to program and drive collective migration in these tissues using bioelectric stimulation. Here, we delivered a unidirectional electrotactic cue using a modified version of our SCHEEPDOG electro-bioreactor (Methods). Briefly, the SCHEEPDOG platform integrates a microfluidic bioreactor containing programmable electrodes around pre-grown tissue arrays. Here, we applied an electric field of 2V/cm for 8h across keratinocyte monolayers patterned and cultured as described previously (Fig. 1F). While all tissues responded strongly the applied field, the nature of the response heavily depended on the collective strength of the tissue (Fig. 1G, Movie S2).

Specifically, changes in collective strength impacted the spatiotemporal response of the tissue with respect to migration speed and directedness (Fig. 2A). While cells in all tissues increased their overall speed during electrotaxis as seen in previous work^6,20,21,34,39,40^, the relative increase in speed varied inversely with collective strength, with weakly collective monolayers migrating at almost twice the speed of strongly collective monolayers under the same electrical stimulation (Figs. 2B, 2C). Faster motion in less strongly collective tissues was consistent with the baseline motility data without stimulation. Although the overall directedness of collective migration during electrotaxis was independent of collective strength, we noted that stronger collectives took longer to align than did weakly collective tissues, with the most strongly collective tissues taking ∼35 minutes longer to align than the other conditions (Figs. 2D, 2E). This clearly demonstrates a competition between the endogenous collective behavior of a tissue and the imposed command, making more strongly collective tissues less responsive to bioelectric cues.

**Figure 2.**
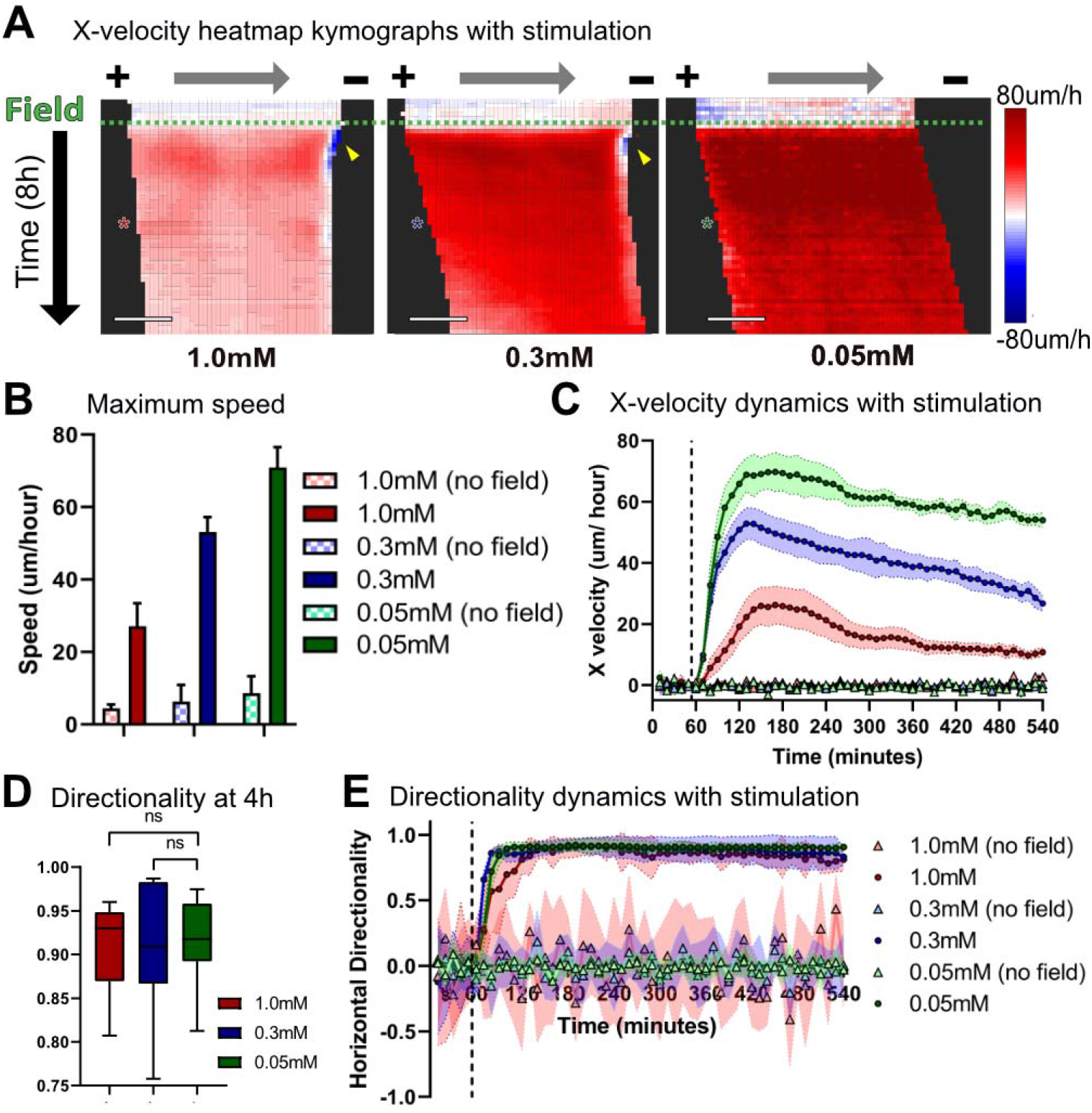
Increased coordination reduces monolayers’ responsivity to electrical stimulation. A) X-velocity heatmap kymograph for 1h control and 8h stimulation. Each square corresponds to 40-45um of the monolayer. Electrical stimulation starts at the green dashed line. Asterisks indicate 4h into electrical stimulation. 10 min/ row. Scale bar = 500um. B) Maximum migration speed for monolayers with and without electrical stimulation. C) Averaged X-velocity of migrating monolayers throughout 1h control (no field) and 8h stimulation. Error bars represent standard deviation across tissues. Dashed vertical lines denote when the field was switched on. Legends identical to B). D) Horizontal directionality at 4h into stimulation. E) Horizontal directionality throughout 1h control and 8h stimulation. Error bars represent standard deviation across tissues. Dashed vertical lines denote when the field was switched on. P values are calculated using unpaired nonparametric Mann-Whitney test with n = 12-15 for each condition.

### Naive collective control can result in catastrophic damage to the tissue

Beyond differences in speed and response time, we observed a far more striking and detrimental phenotype: both our moderately and strongly collective tissues experienced powerful retraction and collapse of their leading edges, with the effect being more pronounced in strongly collective tissues (Figs. 1G, 2A, 3A). Quantifying the dynamics of retraction revealed retraction occurred within 15 minutes of electrical stimulation (Figs. 2A, S3) in the moderate and strong collectives, while weakly collective tissues advanced with no apparent problems. Retraction also caused high cytotoxicity, and a marker for membrane damage (ethidium homodimer, Methods) revealed strong and localized damage all along the retracting edge (Figs. 3A, S4; Movie S3). We quantified the overall effect of retraction by analyzing total leading edge displacement over 8 h of stimulation (Fig. 3B), where we see that strongly collective tissues experienced net negative forward motion, moderately collective tissues recovered some forward motion, and weakly collective tissues advanced nearly 4X over their unstimulated control case.

**Figure 3.**
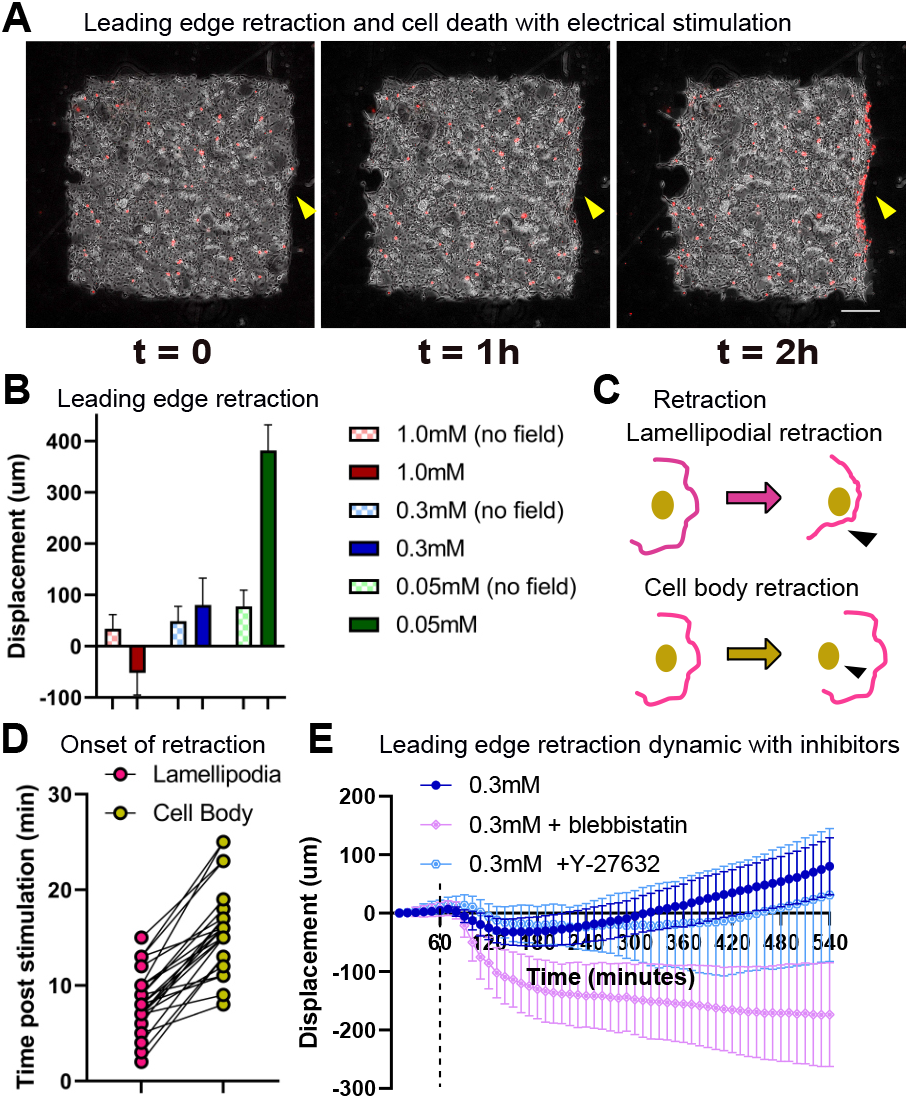
Leading edge retraction and cellular damage with stimulation in highly coordinated monolayers. A) Phase (grey) and EthD-1 dye (red) images throughout 2h electrical stimulation of medium calcium monolayer. Yellow arrows point to cell death and retraction at leading edge. Scale bar = 500um. B) Leading edge displacement after 9h for monolayers with and without electrical stimulation. Error bars represent standard deviation across tissues. C) Schematic of lamellipodial retraction vs. cell body retraction with electrical stimulation. D) Onset time of lamellipodial retraction and cell body retraction post electric stimulation (n = 24). E) Leading edge displacement plot for medium calcium monolayers treated with blebbistatin (light pink) and Y-27632 (light blue) and electrically stimulated. Error bars represent standard deviation across tissues (n = 10). Dashed vertical lines denote when the field was switched on.

To better understand retraction, we analyzed higher frame-rate videos of the process and found that, in all cases, lamellipodial detachment preceded both cell blebbing and eventual retraction of the cell body (Figs. 3C, 3D; Movie S3). Such retraction is strikingly reminiscent of tissue dewetting, a phenomenon in which cellular monolayers detach from the substrate and retract inwards as E-cadherin junctions trigger myosin phosphorylation, increasing cortical tension within the monolayer^41,42^. That we do not observe retraction in single cells at any calcium level is also consistent with dewetting (Movie S4). As dewetting could be delayed by reducing contractility^41^, we hypothesized that disrupting contractility in monolayers would also mitigate leading edge retraction. We used inhibitors to disrupt contractility in electrotaxing cell collectives, by treating monolayers with either blebbistatin or Y-27632 at 20uM for 1h before electrical stimulation^39,43^ and maintaining inhibitor levels during perfusion. However, both inhibitors failed to mitigate retraction—while Y-27632 had little effect, blebbistatin significantly worsened the phenotype (Fig. 3E, Movie S5). This suggests that simple contractility is unlikely to be the dominant driving force in leading edge retraction.

### Cell-cell adhesion is uniquely responsible for bioelectric collective migration control

Based on our data showing a correlation between collective strength and junctional E-cadherin, we hypothesized that E-cadherin-mediated cell-cell adhesion was a likely regulator of electrotactic control. To validate this and to rule out effects from calcium signaling^44–46^, we treated tissues with a known blocking antibody against extracellular E-cadherin (DECMA-1) that specifically targeted and weakened cell-cell adhesion without altering calcium (Fig. 4A)^47^. Addition of the E-cadherin blocking antibody potentially increased unstimulated migration speed within the monolayers at all calcium levels, and significantly reduced overall migration coordination even in the moderate and high calcium samples (Figs. 4B, 4C).

**Figure 4.**
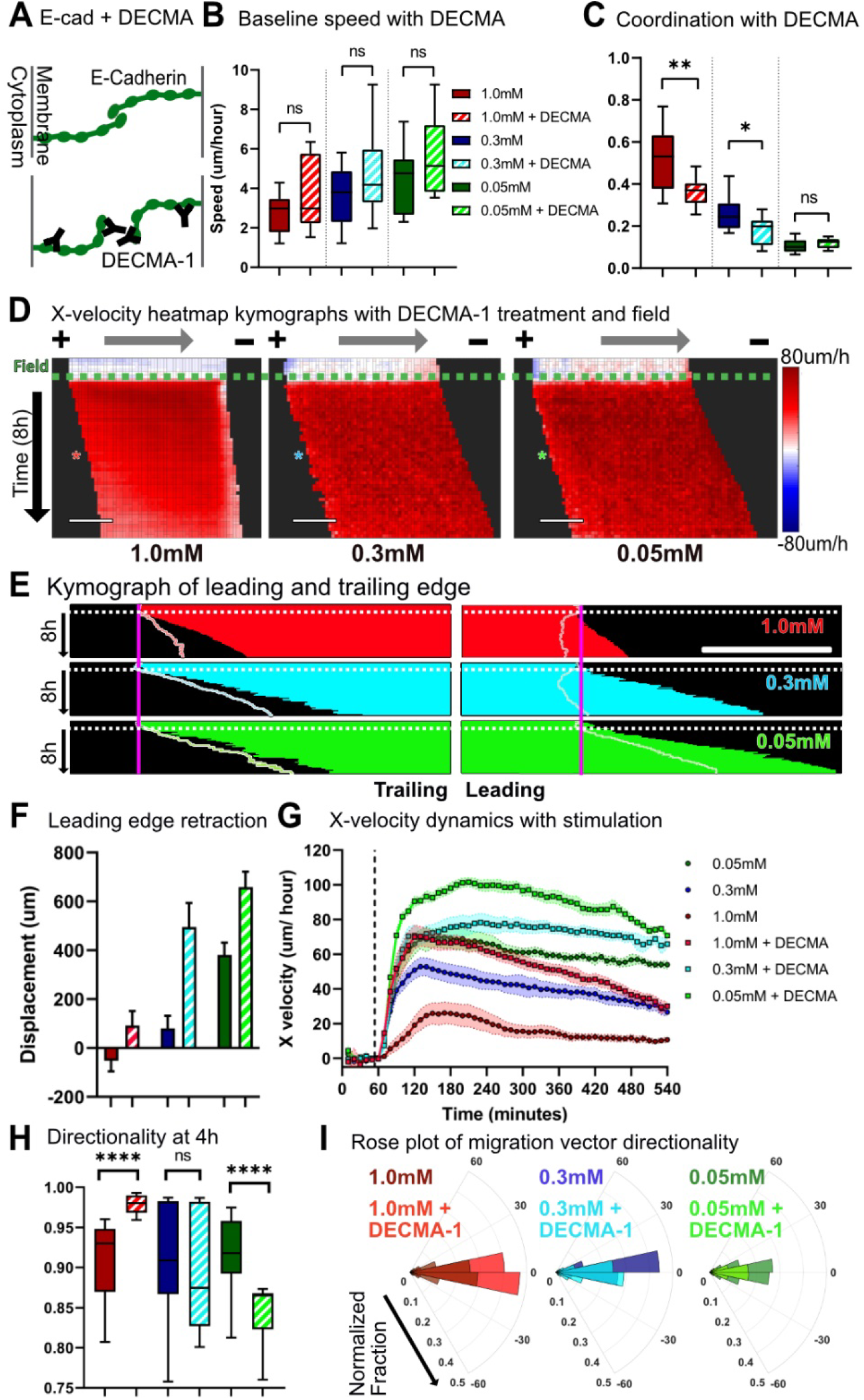
Disrupting E-cadherin junction formation with DECMA-1 reduces coordination and increases controllability. A) Schematic of normal E-cadherin junction formation vs. with DECMA-1 disruption. B) Baseline migration speed for monolayers cultured in varying calcium, with and without DECMA-1. C) Coordination values for monolayers cultured in varying calcium, with and without DECMA-1. Legends identical to B). D) X-velocity heatmap kymograph for monolayers pretreated with DECMA-1 throughout 1h control and 8h stimulation. Each square corresponds to 40-45um of the monolayer. Electrical stimulation starts at the green dashed line. Asterisks indicate 4h into electrical stimulation. 10 min/ row. Scale bar = 500um. E) Kymographs of monolayers pretreated with DECMA-1 throughout 1h control and 8h stimulation. Electrical stimulation starts at white dotted line. Pastel outlines indicate the edge of stimulated monolayers without DECMA-1 at same calcium level. Scale bar = 500um. F) Leading edge displacement after 1h control and 8h stimulation for monolayers with and without DECMA-1 at varying calcium. Legends identical to B). G) X-velocity throughout 1h control and 8h stimulation for monolayers with and without DECMA-1. Error bars represent standard deviation across tissues. Dashed vertical lines denote when the field was switched on. H) Horizontal directionality at 4h into stimulation for monolayers with and without DECMA-1 with varying calcium. I) Polar distribution plot of the velocity vector angle with respect to direction of electrical field. Legends identical to B). P values are calculated using unpaired nonparametric Mann-Whitney test with n = 12-15 for each condition. * corresponds to p < 0.05, ** to p < 0.01, and **** to p < 0.0001.

Having downregulated collective strength of tissues at all three calcium levels, we then tested how they responded to electrical stimulation. DECMA-1 treatment ‘rescued’ forward motion by alleviating retraction in all calcium conditions (Fig. 4D-F, Movie S6). Notably, all tissues experienced improvements to both forward motion (Fig. 4F) and average speed (Fig. 4G). That DECMA treatment improved performance in even low calcium tissues was notable as it implied that even the weak cell-cell adhesion still present in those tissues constrained the electrotactic response. Interestingly, while the overall speed and displacement of tissues were improved by blocking cell-cell adhesion, the accuracy, or directionality of the collective migration response was more nuanced (Fig. 4H). DECMA-1 significantly increased the directionality in strongly collective monolayers while reducing directionality in weakly collective monolayers. To better relate this to accuracy or ‘spread’, we plotted polar histograms of the angles between cell velocity vectors and the electric field vector (Fig. 4I). Specifically, DECMA-1 decreased scattering of electrotactic collective migration in strongly collective monolayers, while treating weakly collective monolayers with DECMA-1 increased scattering in the direction perpendicular to the electrical field making the control less precise (Fig. 4I, right). These data both suggested that overly strong native coordination, mediated specifically by E-cadherin here, can reduce controllability or cause adverse effects such as retraction.

### Disassembly, collective transport, and reassembly of a tissue as a control strategy

Knowing both that strong cell-cell adhesion can limit electrotactic control in skin, and yet E-cadherin is essential for skin function and barrier formation, we sought to develop a more general stimulation strategy to allow us to transiently disrupt cell-cell junctions, use electrotaxis to reshape or move the more susceptible tissue, and then reassemble junctions when the tissue had reached its target location. While DECMA-1 treatment was effective at revealing the role of E-cadherin, it has three significant limitations as a general approach: (1) antibodies are expensive; (2) it is difficult to control how long it will block junctions; and (3) antibodies appear to have a difficult time penetrating very strong cell-cell junctions (Figs. 4D-F, Fig. S5), thereby limiting their overall value in the very tissues we are trying to control more effectively. As an alternative, we tested brief exposure to BAPTA, an extracellular calcium-specific chelator (Methods), and examined how it disrupted E-cadherin junctions in pre-established tissues^48^. Fluorescence imaging of GFP E-cadherin keratinocytes confirmed that 1h of BAPTA treatment applied to tissues with strong E-cadherin junctions could transiently reduce junctional E-cadherin and reduce coordination (Fig. 5A, B).

**Figure 5.**
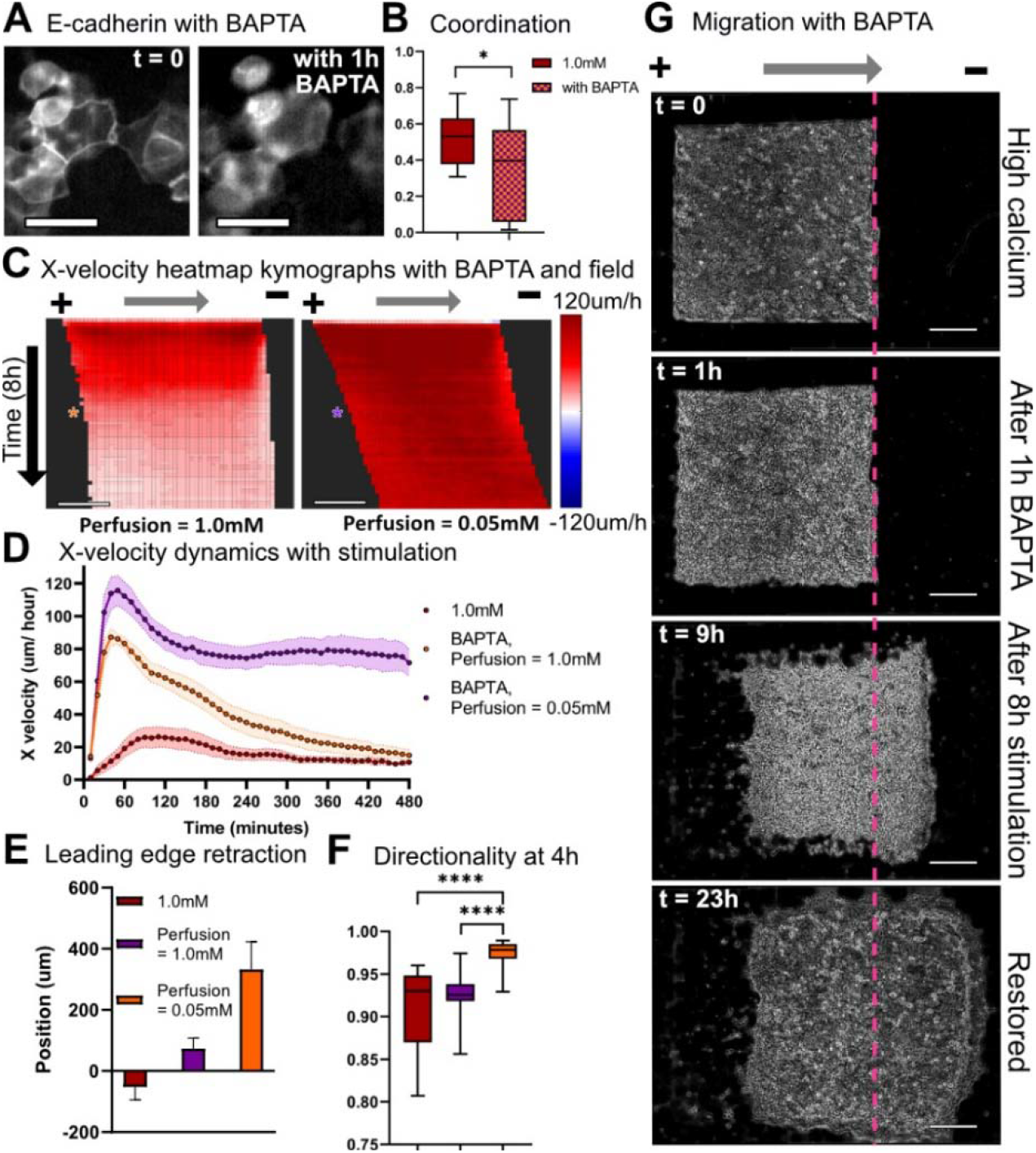
Controllability of highly coordinated monolayers can easily and quickly rescued by acutely altering E-cadherin junctions. A) GFP E-cadherin keratinocyte fluorescence images at t = 0 (left) and with 1h BAPTA treatment (right). Scale bar = 20um. B) Coordination values for high calcium monolayers and high calcium monolayers treated for 1h with 20uM BAPTA. C) X-velocity heatmap kymograph for BAPTA-treated high calcium monolayers stimulated in high and low calcium media. Asterisks indicate 4h into electrical stimulation. 10 min/ row. Scale bar = 500um. D) X-velocity throughout 8h stimulation for high calcium monolayers and high calcium monolayers treated with BAPTA and stimulated in high or low calcium media. Error bars represent standard deviation across tissues. E) Leading edge displacement of BAPTA treated high calcium monolayers after 8h stimulation in high and low calcium media. Error bars represent standard deviation across tissues. F) Horizontal directionality at 4h into stimulation. G) Phase image of high calcium keratinocyte monolayers at t = 0, treated 1h with BAPTA, electrically stimulated in low calcium media for 8h, and restored in high calcium media for 14h. Image at t = 0, t = 1h after 1h BAPTA treatment, t = 9h after 8h stimulation in low calcium media, and t = 17h after 14h restoration in high calcium media. Scale bar = 500um. P values are calculated using unpaired nonparametric Mann-Whitney test with n = 12-15 for each condition. ∗ corresponds to p < 0.05, and ∗∗∗∗ to p < 0.0001.

To test how rapid chelation affected the controllability of strongly collective monolayers, we treated monolayers with BAPTA for 1h, washed out the chelator, and returned the monolayers to BAPTA-free, high calcium media for electrical stimulation. 1h of BAPTA treatment boosted controllability in strongly collective monolayers, with treated monolayers exhibiting both significantly increased migration speed and reduced leading edge retraction (Fig. 5B). However, these benefits were short-lived and speed and displacement drastically decreased over time (Figs. 5C-E, ‘orange’) likely as cell-cell junctions eventually re-engaged due to the high calcium concentration (Fig. S6). To prevent the gradual restoration of junctions, we maintained tissues in low calcium media after washing out BAPTA. These tuned tissues and exhibited a nearly 5X increase in maximum speed, strong leading edge displacement, and high alignment with the field command (Figs. 5B-E, ‘purple’).

Having confirmed that transient chelation could dramatically increase controllability, we then examined if we could restore the monolayer to its initial, highly coordinated state by removing the electrical field and returning disrupted monolayers to high calcium media, allowing the calcium to reestablish junctions. E-cadherin fluorescence imaging shows that disrupted monolayers returned to high calcium media overnight regained their contact with neighbors and reestablished strong E-cadherin junctions (Fig. S7). Timelapse imaging of the entire process—BAPTA treatment of strongly collective monolayers, migration in low calcium media, and restoration in high calcium media—demonstrates how a difficult to control tissue can be transformed to a more susceptible tissue, maneuvered to a desired location an arbitrary distance away, and then reassembled (Figure 5I, Movie S8). In this case, while we do still note a thin zone of membrane damage at the initial leading edge (Movie S8, red band at the rightward edge), this no longer causes retraction and the tissue instead surges forward as a cohesive unit.

### Accelerating bioelectric healing *in vitro* by manipulating the strength of cell-cell adhesion

Combining pharmacological perturbations with bioelectric cues to improve tissue response suggests practical avenues to engineering the behavior of otherwise recalcitrant tissues for practical purposes. To demonstrate this, we attempted to electrically accelerate *in vitro* wound healing of a strongly collective skin layer. In this case, naïve stimulation would trigger a collapse or at best no edge outgrowth (Figs. 2-3), but the disassembly/reassembly process described above should enable complete, expedited healing. To test this, we created a wound gap across a strongly collective, high-calcium skin layer and then reconfigured the electrodes in SCHEEPDOG to generate an electric field that converged on the middle of the wound to drive each side of the tissue inwards^49^ (Methods). Identical to the scheme described above, strongly collective monolayers were treated with BAPTA for 1h, stimulated in low calcium media for 12h, and restored in high calcium media. The increase of wound closure rate for BAPTA + electrically stimulated tissues compared non-stimulated strongly collective monolayers is clearly visible in the timelapse panels (Fig. 6, Movie S9). Monolayers moved towards each other rapidly during the 12h stimulation and successfully merged soon after they were returned to high calcium media to restore their initial state. These data demonstrate both how controllability of tissues can be dynamically tuned, and how such tuning can be used to practical effect—in this case, increasing the baseline wound closure rate by ∼2.5X.

**Figure 6.**
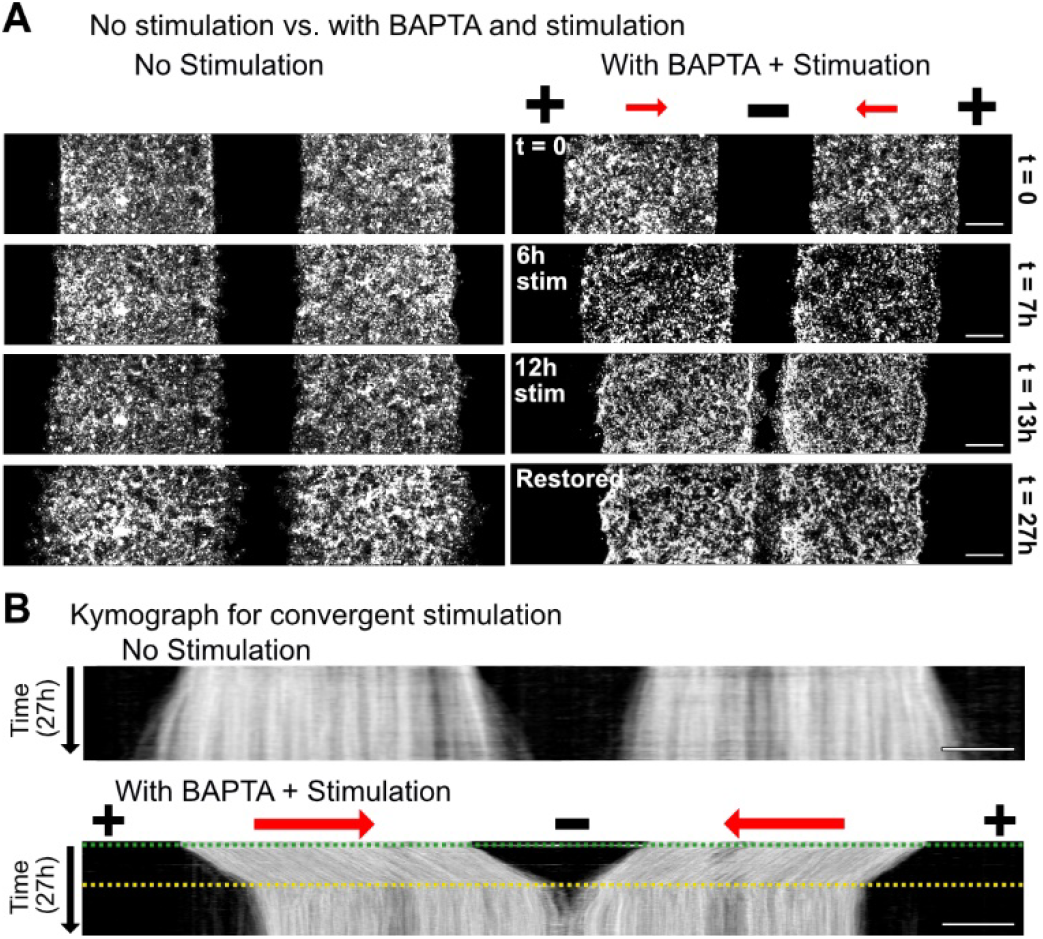
Accelerated wound healing using electrical stimulation and manipulation of cell-cell adhesion strengths. A) Wound closure: fluorescence images for unstimulated high calcium monolayers (left) and high calcium monolayer treated with BAPTA, convergently stimulated in low calcium media for 12h, and incubated in high calcium media for 14h (right). Scale bar = 500um. B) Kymograph of unstimulated high calcium monolayer (top) and high calcium monolayer treated with BAPTA, convergently stimulated in low calcium media for 12h, and incubated in high calcium media for 14h (bottom). Green dashed line indicates when the stimulation was switched on and media was changed to low calcium media, and yellow dashed line indicates when the stimulation was switched off and monolayers were returned to high calcium media. Scale bar = 500um.

### Discussion: “If you can’t join it, then beat it.”

Our work demonstrates that the more strongly collective a given tissue is—determined here by cell-cell adhesion and native coordination levels—the more difficult it may be to externally program the behavior of that tissue as the command and the native behaviors compete with each other. A corollary to this is that, rather than synergizing with an existing collective behavior it can be beneficial to weaken, override, or ‘beat it’. In particular, our results demonstrate that we can better optimize the ‘controllability’ of a cellular collective by both applying an appropriate external stimulus, and also modifying the internal, collective imperatives of the target system to mitigate the chance of conflict between imperatives.

Surprisingly, the consequences of ignoring the potential conflict between the command and natural imperative of a tissue can be quite drastic. While programmed electrotaxis of layers of weakly coupled primary mouse skin cells allowed for clean, large scale control over tissue migration, the same electrical stimulation applied to strongly collective skin layers resulted in not only collapse of the leading edge of the tissue, but also considerable membrane damage in those cells at the leading edge (Figs. 2,3). Some level of supracellular differences in behavior across an electrotaxing tissue—where the edges of a tissue seem less responsive than the bulk—have been noted in several prior electrotaxis studies in different models^6,21,34^, but the collapse we see here has not been previously reported. Further, that inhibiting cell contractility (Fig. 3) worsened the problem here suggests that collective contractility is not to blame for sub-optimal electrotaxis and is consistent with prior data indicating that inhibiting myosin-mediated contractility does not abolish collective electrotaxis^39^. Further work on actual cytoskeletal morphology and behavior at the leading edge of driven, collectively migrating tissues seem necessary to better clarify the role of the cytoskeleton in the collapse we observe.

However, we were able to completely mitigate edge collapse and restore sustained directed motion across a whole tissue by specifically targeting E-cadherin to weaken cell-cell adhesion strength. Cell-cell adhesion, often regulated by E-cadherin, plays a critical role in collective cell migration as cell-cell junctions allow intimate coupling of physical forces and mechanical signaling across cells, which can enable long-range coordination and the emergence of collective motion^50,51^. Our data linking reduced E-cadherin levels to weaker baseline coordination (Figs. 1B-D, 4C), and the results of specific inhibition of E-cadherin junctions (Fig. 4D-I) support the concept that targeting E-cadherin tipped the balance in favor of electrotaxis, allowing the electrical cue to outcompete the now weaker internal collective prerogatives of the tissue. When the results are considered alongside prior findings where E-cadherin knock-down diminished electrotaxis in immortalized epithelial cells^8,52^, despite the complications in direct comparison due to differences in the cell type and baseline collective behaviors, the emerging story shows that while E-cadherin appears to be play a major role in regulating collective electrotaxis, either too little or too much cell-cell adhesion can detrimentally affect controllability. Hence, there appears to be a ‘goldilocks’ window for cell-cell adhesion strength and effective electrotactic control, and native cell coordination should be treated as an independent variable to be modified as needed to optimize controllability, such as with electrotaxis.

This ability to independently tune internal collective strength and externally electrically stimulate a tissue suggested a solution to the problem of controlling strongly collective tissues: (1) transiently weaken internal collective coupling in a tissue; (2) bioelectrically drive the more controllable tissue to a target location or configuration; and (3) fully restore cell-cell coupling and tissue integrity at the new location. This approach ultimately allowed us to accelerate the collective healing process of a strongly collective, injured skin layer such that it healed at least twice as quickly as the control. Unexpectedly, we noted that electrotactic performance during this process of dynamically adjusting collective strength was improved, in terms of both speed and directionality, compared to tissues that began as weak collectives (Fig. 2 versus Fig. 5). That we can not only control collective cell behaviors, but also begin to optimize this control is exciting as there has been tremendous recent effort towards developing bioelectric wound dressings capable of improving healing *in vivo*^53–56^. We hope our results and control paradigms here might help enable next-generation biointerfaces for clinical applications, *a process that has been stalled despite promising results as the underlying mechanisms are difficult to characterize and observe, and there are few formal ‘design rules’ for thinking about how to improve performance*^57^

More broadly, our findings highlight underlying fundamental principles across collective systems and are in line with diverse examples of collective motion and control. For example, swarm theory predicts that overly strong collective coupling can reduce the responsiveness of the system to external perturbations, a finding consistent with experimental data across multiple systems^58^. Panic in human groups can increase the strength and distance of correlated motion within the group, inhibiting the group’s ability to efficiently take advantage of exit cues and doorways during escape conditions^59^. Similarly, swarms of locust nymphs have been shown to be more difficult to redirect the denser and more aligned the natural structure of the swarm is^60,61^. Finally, penguin huddles exhibit a natural clustering tendency, creating a jamming transition that would cause penguins on the outside of the group to die of exposure unless penguin clusters break symmetry and push their neighbors to transiently fluidize this jammed state and allow circulation from the outside in^62^. In each of these examples, the underlying collective behaviors define the properties of the group, with stronger collectivity and coordination reducing the responsiveness and controllability of collectives. Given key similarities across collective systems, it is likely that there are many more guidelines from natural collective processes that we can take inspiration from to improve our ability to program tissues.

## References

1. Friedl, P. & Gilmour, D. Collective cell migration in morphogenesis, regeneration and cancer. Nat. Rev. Mol. cell Biol. 10, 445 (2009).

2. Li, J. & Lin, F. Microfluidic devices for studying chemotaxis and electrotaxis. Trends Cell Biol. 21, 489–497 (2011).

3. Berthier, E. & Beebe, D. J. Gradient generation platforms: new directions for an established microfluidic technology. Lab Chip 14, 3241–3247 (2014).

4. Weitzman, M. & Hahn, K. M. Optogenetic approaches to cell migration and beyond. Curr. Opin. Cell Biol. 30, 112–120 (2014).

5. Xiong, S., Gao, H., Qin, L., Jia, Y.-G. & Ren, L. Engineering topography: Effects on corneal cell behavior and integration into corneal tissue engineering. Bioact. Mater. 4, 293–302 (2019).

6. Zajdel, T. J., Shim, G., Wang, L., Rossello-Martinez, A. & Cohen, D. J. SCHEEPDOG: Programming Electric Cues to Dynamically Herd Large-Scale Cell Migration. Cell Syst. 10, 506-514.e3 (2020).

7. Gokoffski, K. K., Jia, X., Shvarts, D., Xia, G. & Zhao, M. Physiologic Electrical Fields Direct Retinal Ganglion Cell Axon Growth In Vitro. Invest. Ophthalmol. Vis. Sci. 60, 3659–3668 (2019).

8. Li, L. et al. E-cadherin plays an essential role in collective directional migration of large epithelial sheets. Cell. Mol. Life Sci. 69, 2779–2789 (2012).

9. Mayor, R. & Etienne-Manneville, S. The front and rear of collective cell migration. Nature Reviews Molecular Cell Biology (2016). doi:10.1038/nrm.2015.14

10. Petridou, N. I. & Heisenberg, C. Tissue rheology in embryonic organization. EMBO J. 38, 1–13 (2019).

11. Tlili, S. et al. Collective cell migration without proliferation: Density determines cell velocity and wave velocity. R. Soc. Open Sci. 5, (2018).

12. Barriga, E. H., Franze, K., Charras, G. & Mayor, R. Tissue stiffening coordinates morphogenesis by triggering collective cell migration in vivo. Nature 554, 523–527 (2018).

13. Kuriyama, S. et al. In vivo collective cell migration requires an LPAR2-dependent increase in tissue fluidity. J. Cell Biol. 206, 113–127 (2014).

14. Vannier, C., Mock, K., Brabletz, T. & Driever, W. Zeb1 regulates E-cadherin and Epcam (epithelial cell adhesion molecule) expression to control cell behavior in early zebrafish development. J. Biol. Chem. 288, 18643–18659 (2013).

15. McCaig, C. D., Song, B. & Rajnicek, A. M. Electrical dimensions in cell science. J. Cell Sci. (2009). doi:10.1242/jcs.023564

16. Cortese, B., Palama, I. E., D’Amone, S. & Gigli, G. Influence of electrotaxis on cell behaviour. Integr. Biol. 6, 817–830 (2014).

17. Nuccitelli, R. A role for endogenous electric fields in wound healing. Curr. Top. Dev. Biol. 58, 1–26 (2003).

18. Allen, G. M., Mogilner, A. & Theriot, J. A. Electrophoresis of cellular membrane components creates the directional cue guiding keratocyte galvanotaxis. Curr. Biol. 23, 560–568 (2013).

19. Sprott, D., Aceh, kue tradisional khas & Sprott, D. PI3K inhibition reverses migratory direction of single cells but not cell groups in electric field. Block Caving – A Viable Altern. 21, 1–9 (2020).

20. Lalli, M. L. & Asthagiri, A. R. Collective Migration Exhibits Greater Sensitivity But Slower Dynamics of Alignment to Applied Electric Fields. Cell. Mol. Bioeng. 8, 247–257 (2015).

21. Cho, Y., Son, M., Jeong, H. & Shin, J. H. Electric field-induced migration and intercellular stress alignment in a collective epithelial monolayer. Mol. Biol. Cell 29, 2292–2302 (2018).

22. Guo, X. et al. The galvanotactic migration of keratinocytes is enhanced by hypoxic preconditioning. file:///C:/Users/gawoo/Downloads/Papers/nihms-415862.pdfScientific Reports 5, 1–13 (2015).

23. Zhao, M. Electrical fields in wound healing-An overriding signal that directs cell migration. Semin. Cell Dev. Biol. 20, 674–682 (2009).

24. Elias, P. M., Ahn, S. K., Brown, B. E., Crumrine, D. & Feingold, K. R. Origin of the epidermal calcium gradient: Regulation by barrier status and role of active vs passive mechanisms. J. Invest. Dermatol. (2002). doi:10.1046/j.1523-1747.2002.19622.x

25. Menon, G. K., Grayson, S. & Elias, P. M. Ionic calcium reservoirs in mammalian epidermis: Ultrastructural localization by ion-capture cytochemistry. J. Invest. Dermatol. 84, 508–512 (1985).

26. Menon, G. K., Elias, P. M., Lee, S. H. & Feingold, K. R. Localization of calcium in murine epidermis following disruption and repair of the permeability barrier. Cell Tissue Res. (1992). doi:10.1007/BF00645052

27. Yuspa, S. H., Kilkenny, A. E., Steinert, P. M. & Roop, D. R. Expression of murine epidermal differentiation markers is tightly regulated by restricted extracellular calcium concentrations in vitro. J. Cell Biol. 109, 1207–1217 (1989).

28. Wang, F., Chen, S., Liu, H. B., Parent, C. A. & Coulombe, P. A. Keratin 6 regulates collective keratinocyte migration by altering cell–cell and cell–matrix adhesion. J. Cell Biol. 217, 4314–4330 (2018).

29. Vasioukhin, V. & Fuchs, E. Actin dynamics and cell-cell adhesion in epithelia. Curr. Opin. Cell Biol. 13, 76–84 (2001).

30. Vedula, S. R. K. et al. Emerging modes of collective cell migration induced by geometrical constraints. Proc. Natl. Acad. Sci. U. S. A. 109, 12974–12979 (2012).

31. Benjamin, J. M. et al. αE-catenin regulates actin dynamics independently of cadherin-mediated cell-cell adhesion. J. Cell Biol. 189, 339–352 (2010).

32. Xi, W., Sonam, S., Beng Saw, T., Ladoux, B. & Teck Lim, C. Emergent patterns of collective cell migration under tubular confinement. Nat. Commun. 8, (2017).

33. Petitjean, L. et al. Velocity fields in a collectively migrating epithelium. Biophys. J. 98, 1790–1800 (2010).

34. Cohen, D. J., Nelson, W. J. & Maharbiz, M. M. Galvanotactic control of collective cell migration in epithelial monolayers. Nat. Mater. 13, 409–417 (2014).

35. Vaezi. Actin Cable Dynamics and Rho/Rock Orchestrate a Polarized Cytoskeletal Architecture in the Early Steps of Assembling a Stratified Epithelium. J. Appl. Econ. Sci. 4, 169–184 (2009).

36. Vasioukhin, V., Bauer, C., Yin, M. & Fuchs, E. Directed Actin Polymerization Is the Driving Force for Epithelial Cell-Cell Adhesion-catenin associates with several other actin-binding. AJs (Drubin Nelson 100, 209–219 (2000).

37. Cavagna, A. et al. Scale-free correlations in starling flocks. Proc. Natl. Acad. Sci. U. S. A. 107, 11865–11870 (2010).

38. Spatarelu, C.-P. et al. Biomechanics of Collective Cell Migration in Cancer Progression: Experimental and Computational Methods. ACS Biomater. Sci. Eng. 5, 3766–3787 (2019).

39. Bashirzadeh, Y., Poole, J., Qian, S. & Maruthamuthu, V. Effect of pharmacological modulation of actin and myosin on collective cell electrotaxis. Bioelectromagnetics 39, 289–298 (2018).

40. Zhao, M., Agius-Fernandez, A., Forrester, J. V. & McCaig, C. D. Directed migration of corneal epithelial sheets in physiological electric fields. Investig. Ophthalmol. Vis. Sci. 37, 2548–2558 (1996).

41. Pérez-González, C. et al. Active wetting of epithelial tissues. Nat. Phys. 15, 79–88 (2019).

42. Douezana, S. & Brochard-Wyart, F. Dewetting of cellular monolayers. Eur. Phys. J. E 35, 0–5 (2012).

43. Saltukoglu, D. et al. Spontaneous and electric feld-controlled front-rear polarization of human keratinocytes. Mol. Biol. Cell 26, 4373–4386 (2015).

44. Mycielska, M. E. & Djamgoz, M. B. A. Cellular mechanisms of direct-current electric field effects: Galvanotaxis and metastatic disease. J. Cell Sci. 117, 1631–1639 (2004).

45. Shanley, L. J., Walczysko, P., Bain, M., MacEwan, D. J. & Zhao, M. Influx of extracellular Ca2+ is necessary for electrotaxis in Dictyostelium. J. Cell Sci. 119, 4741–4748 (2006).

46. Bikle, D. D., Ratnam, A., Mauro, T., Harris, J. & Pillai, S. Changes in calcium responsiveness and handling during keratinocyte differentiation: Potential role of the calcium receptor. J. Clin. Invest. 97, 1085–1093 (1996).

47. Perez, T. D. & Nelson, W. J. Cadherin Adhesion: Mechanisms and Molecular Interactions. 5, 3–21 (2004).

48. Kim, S. A., Tai, C. Y., Mok, L. P., Mosser, E. A. & Schuman, E. M. Calcium-dependent dynamics of cadherin interactions at cell-cell junctions. Proc. Natl. Acad. Sci. U. S. A. 108, 9857–9862 (2011).

49. Zajdel, T. J., Shim, G. & Cohen, D. J. Come together: bioelectric healing-on-a-chip. bioRxiv (2020). doi:10.1101/2020.12.29.424578

50. Shellard, A. & Mayor, R. Supracellular migration - Beyond collective cell migration. Journal of Cell Science (2019). doi:10.1242/jcs.226142

51. De Pascalis, C. & Etienne-Manneville, S. Single and collective cell migration: The mechanics of adhesions. Mol. Biol. Cell 28, 1833–1846 (2017).

52. Lalli, M. L., Wojeski, B. & Asthagiri, A. R. Label-Free Automated Cell Tracking: Analysis of the Role of E-cadherin Expression in Collective Electrotaxis. Cell. Mol. Bioeng. 10, 89–101 (2017).

53. Long, Y. et al. Effective Wound Healing Enabled by Discrete Alternative Electric Fields from Wearable Nanogenerators. ACS Nano 12, 12533–12540 (2018).

54. Boateng, J. & Catanzano, O. Advanced Therapeutic Dressings for Effective Wound Healing - A Review. J. Pharm. Sci. 104, 3653–3680 (2015).

55. Li, M. et al. Toward Controlled Electrical Stimulation for Wound Healing Based on a Precision Layered Skin Model. ACS Appl. Bio Mater. 3, 8901–8910 (2020).

56. Zhao, Z. et al. Optimization of Electrical Stimulation for Safe and Effective Guidance of Human Cells. Bioelectricity 2, 372–381 (2020).

57. Zhao, M., Penninger, J. & Isseroff, R. R. Electrical Activation of Wound-Healing Pathways. Adv. Skin Wound Care 1, 567–573 (2010).

58. Bouffanais, R. A Computational Approach to Collective Behaviors. in Design and Control of Swarm Dynamics 95–104 (Springer Singapore, 2016). doi:10.1007/978-981-287-751-2_6

59. Jiang, L., Li, J., Shen, C., Yang, S. & Han, Z. Obstacle optimization for panic flow - Reducing the tangential momentum increases the escape speed. PLoS One 9, 21–25 (2014).

60. Buhl, J. et al. From Disorder to Order in Marching Locusts. Science (80-.). 312, 1402–1406 (2006).

61. Yates, C. A. et al. Inherent noise can facilitate coherence in collective swarm motion. Proc. Natl. Acad. Sci. U. S. A. 106, 5464–5469 (2009).

62. Zitterbart, D. P., Wienecke, B., Butler, J. P. & Fabry, B. Coordinated movements prevent jamming in an emperor penguin huddle. PLoS One 6, 5–7 (2011).

