## Supplementary material for "Cellular crowd control: overriding endogenous cell coordination makes cell migration more susceptible to external programming": Methods

**Cell Culture**

*Cell maintenance*

Primary keratinocytes and GFP E-cadherin keratinocytes were harvested from mice (courtesy of the Devenport Laboratory, Princeton University). Cells were cultured in E-medium supplemented with 15% serum and 50 µM calcium^1^. All cells were maintained at 37 °C under 5% CO2 and 95% relative humidity. Cells were split before 70% confluence and passage number was kept above 35 for all experiments. A 200mM Ca2+ stock (courtesy of the Devenport Laboratory, Princeton University) was diluted in E-Media to adjust the media calcium level as desired. Media was vortexed for 10 s after the addition of calcium and kept overnight at 4C for even mixing.

*Cell patterning*

To seed monolayers in the stimulation zone and control their shapes and sizes, a silicone stencil of 250um thickness (Bisco HT-6240, Stockwell Elastomers) containing five 2x2mm^2^ square microwells was cut and applied to the center of the culture substrate that was treated with fibronectin to provide a matrix for cellular adhesion (protein dissolved to 50 mg/mL in DI water, applied to the dish for 30 min at 37C, then washed three times with DI water). The microwells were spaced 4-6mm from its neighbors to ensure each monolayer had access to sufficient nutrients. With the stencil in place, a seeding solution of cells was prepared at a density of 2.5× 10^6^ cells, counted using Corning Cell Counter (Corning) and 2.5uL of the cell solution was pipetted into the stencils. The cells were allowed to settle for 6 h, and extra humidification media was added to the periphery of the tissue culture dish. Once the cells had adhered to the substrate, sufficient media supplemented with calcium was added to fill the dish. The stencils were removed 14h after incubation for assembly.

*Inhibitor assays*

Monolayers were treated either 20uM blebbistatin (biogems) or Y-27632 (Selleckchem) for 1h before stimulation after the stencils were removed to prevent the inhibitors from being absorbed into the PDMS. Identical concentration of inhibitors was supplemented in the perfusion media to prevent the inhibitors from being washing out during stimulation.

*Live/Dead assay*

Monolayers were treated with 1µM EthD-1 solution (LIVE/DEAD Viability/Cytotoxicity Kit, Invitrogen) for 1h before stimulation after the stencils were removed. Identical concentration of EthD-1 was supplemented in the perfusion media to prevent the reagent washing out during stimulation.

*Cell-cell adhesion disruption*

To disrupt E-cadherin junction formation, cells were first seeded at desired density and allowed to settle for 6h. Calcium supplemented medium containing DECMA-1 at 50µg/mL concentration was added to fill the dish for overnight incubation. After the 14h incubation, DECMA-1 was washed out before the device was assembled for stimulation. For rapid E-cadherin junction disruption, high calcium monolayers were treated with 20µM BAPTA (Tetrasodium salt (cell impermeant), ThermoFisher) for 1h after the 14h incubation. The chelator was washed out directly before the electrical stimulation.

**Electrical field stimulation**

*1D stimulation*

This device assembly is a modified version of the SCHEEPDOG bioreactor from our prior work, and our methods are similar to those published previously^2^. A Kiethly source meter (Kiethly 2450 Textronix) provided current to the silver chloride electrode pairs while an USB oscilloscope (Analog Discovery 2, Digilent Inc.) measured the voltage across the pair of recording electrodes using titanium wires (0.5 mm diameter, Alfa Aesar) as probes. A custom MATLAB script was written to use closed-loop feedback control to adjust the output current from the source meter to maintain the electrical field strength across the microfluidics channel constant at 2V/cm throughout the stimulation period.

*Convergent stimulation*

Device fabrication and instrumentation were executed as published^3^. Two 2x2mm^2^ monolayers spaced 1mm apart with the cathode aligned over the center of the spacing. Anodes were aligned over the outer edges of monolayers to apply convergent electrical stimulation for 12h, with the field strength was kept constant at 2V/cm.

**Microscopy and Imaging**

*Microscopy*

All images were acquired on an automated Zeiss (Observer Z1) inverted fluorescence microscope equipped with an XY motorized stage and controlled using Slidebook (Intelligent Imaging Innovations, 3i). The microscope was fully incubated at 37 °C, and 5% CO2 was constantly bubbled into media reservoir during perfusion. Phase imaging was performed using a 5X/0.16 phase-contrast objective. Fluorescence imaging was performed using a metal halide lamp (xCite 120, EXFO). Immunofluorescence imaging for E-cadherin used a FITC filter set with a 20x/0.8 fluorescence objective and 500 msec exposure. GFP E-cadherin keratinocyte fluorescence imaging used a FITC filter set with a 20x/0.8 fluorescence objective and 200 msec exposure. Fluorescence imaging for EthD-1 used an RFP filter set with a 5X/0.16 phase-contrast objective and 100ms exposure time. Fluorescence imaging for membrane dye used in convergent stimulation used a Cy5 filter set with a 5X/0.16 fluorescent objective and 350ms exposure time. Images were taken at 1 min or 10 min intervals as indicated in the text.

*Immunofluorescence Staining*

Cells were fixed in 4% paraformaldehyde (PFA) solution made by diluting a 16% PFA solution (Pierce 16% Formaldehyde, Thermo Scientific) in PBS. After 10 minutes, the cells were washed with PBS twice then permeabilized with 0.1% Triton X-100 (Sigma Aldrich) solution in PBS for 10 minutes. The cells were washed with PBS-Triton solution twice, then incubated with 1% Bovine Serum Albumin solution (Thermo Fisher) for 15 minutes for blocking for 1h. Primary antibody solution (CD324 (E-Cadherin) Monoclonal Antibody (DECMA-1), eBioscience) diluted to 1:1000 in PBS was added and incubated for 1h. Cells were incubated with BSA solution again for 15 min and washed three times with PBS-Triton solution. The secondary antibody solution (Goat anti-Rabbit IgG (H+L) Alexa Fluor Plus 647, Invitrogen) diluted to 1:1000 in PBS was added and incubated for 30 mins. Cells were washed 3 times in PBS-Triton solution, washed three times in PBS, and maintained in PBS at 4°C. All incubations and washes were done at room temperature.

*Membrane Dye*

Membrane dye (CellBright Red Cytoplasmic Membrane Dye, Biotium) was added to the seeding solution at 1:400 dilution for the convergent stimulation assay. Dye was added at the seeding stage as it could not infiltrate high calcium monolayers if added after the formation of E-cadherin junctions.

**Quantification and statistical analysis**

*Image processing and analysis*

All post-processing of monolayer microscopy images was performed using FIJI^4^. Images were collected sequentially, stitched, and template matched to correct for stage drift prior to being analyzed.

*Particle Image Velocimetry*

Velocity vector fields were generated using PIVLab, a MATLAB plugin performing FFT-based PIV^5^. PIV analysis was performed over the entire monolayer and masked to exclude background noise. Iterative window analysis was performed using first 128x128 pixel windows followed by 64x64 pixel windows, both with 50% step overlap. Vector validation excluded vectors beyond 5 standard deviations and replaced with interpolated vectors. The vector fields were imported into MATLAB to calculate coordination, velocity, speed, and directionality. X-velocity heatmap kymographs were generated by calculating the average velocity at each horizontal point across the entire monolayer at each timepoint, then temporally stacking rows. Velocity correlations with 8 nearest neighbors were calculated for each vector within the monolayer using the following equation^6,7^ where N refers to the total number of vectors being analyzed, n is the number of neighboring objects (as we only look at nearest neighbors, this is 8) from the current object i, and j refers to 8 neighboring objects):

$$Correlation = \frac{1}{N}\sum_{i = 1}^{N} \left( \frac{1}{n}\sum_{j = 1}^{n} \left( \frac{\vec{v_{i}}\cdot\vec{v}_{j}}{\left| \vec{v_{i}} \right|\left| \vec{v_{j}} \right|} \right) \right)$$

*Junctional E-cadherin quantification*

Fluorescence images were tiled using FIJI to create a composite image and was 2x2 binned to reduce noise. 10 junctions were chosen at random per calcium concentration, and a rectangle of 20x100 pixel lengths (approximately 6x30 um) was drawn across each junction, the long edge perpendicular to the junction, and the intensity values were averaged across the short edge using ‘Plot Profile’ plugin in FIJI. The normalized junctional E-cadherin was measured by calculating [Max. Signal – Min. Signal]/ [Min. Signal] of fluorescence intensity across each line.

*Boundary edge displacement kymographs*

Kymographs were produced using FIJI and MATLAB. For the kymographs, the X-position for the leading and trailing edge of the monolayers was averaged for each monolayer using a sum averaging algorithm, then stacked temporally. The kymographs from multiple monolayers were then averaged using MATLAB for each experimental condition. Leading edge displacement was calculated by measuring the distance between the initial averaged X-position of the monolayer before stimulation and the final averaged X-position post-stimulation. For the converging stimulation assay, images were masked to reduce background noise and increased in intensity with MATLAB to compensate for the lack of dye in newly proliferated cells. Masks were created by Gaussian-blurring phase or fluorescence images and thresholding.

*Retraction*

Cell body and lamellipodial retraction at the leading edges were hand-tracked to determine how much time had elapsed since the start of electrical stimulation until the onset of each condition using timelapse videos (1min/ frame) of medium calcium monolayers stimulated at 2V/cm.

1.	Gonzales, K. A. U. & Fuchs, E. Skin and Its Regenerative Powers: An Alliance between Stem Cells and Their Niche. *Dev. Cell* **43**, 387–401 (2017).

2.	Zajdel, T. J., Shim, G., Wang, L., Rossello-Martinez, A. & Cohen, D. J. SCHEEPDOG: Programming Electric Cues to Dynamically Herd Large-Scale Cell Migration. *Cell Syst.* **10**, 506-514.e3 (2020).

3.	Zajdel, T. J., Shim, G. & Cohen, D. J. Come together: bioelectric healing-on-a-chip. *bioRxiv* (2020). doi:10.1101/2020.12.29.424578

4.	Schindelin, J. *et al.* Fiji: An open-source platform for biological-image analysis. *Nat. Methods* **9**, 676–682 (2012).

5.	Thielicke, W. & Stamhuis, E. PIVlab--towards user-friendly, affordable and accurate digital particle image velocimetry in MATLAB. *J. Open Res. Softw.* **2**, (2014).

6.	Angelini, T. E., Hannezo, E., Trepat, X., Fredberg, J. J. & Weitz, D. A. Cell migration driven by cooperative substrate deformation patterns. *Phys. Rev. Lett.* **104**, 168104 (2010).

7.	Haga, H., Irahara, C., Kobayashi, R., Nakagaki, T. & Kawabata, K. Collective movement of epithelial cells on a collagen gel substrate. *Biophys. J.* **88**, 2250–2256 (2005).
