## Supplementary figures for "Cellular crowd control: overriding endogenous cell coordination makes cell migration more susceptible to external programming"

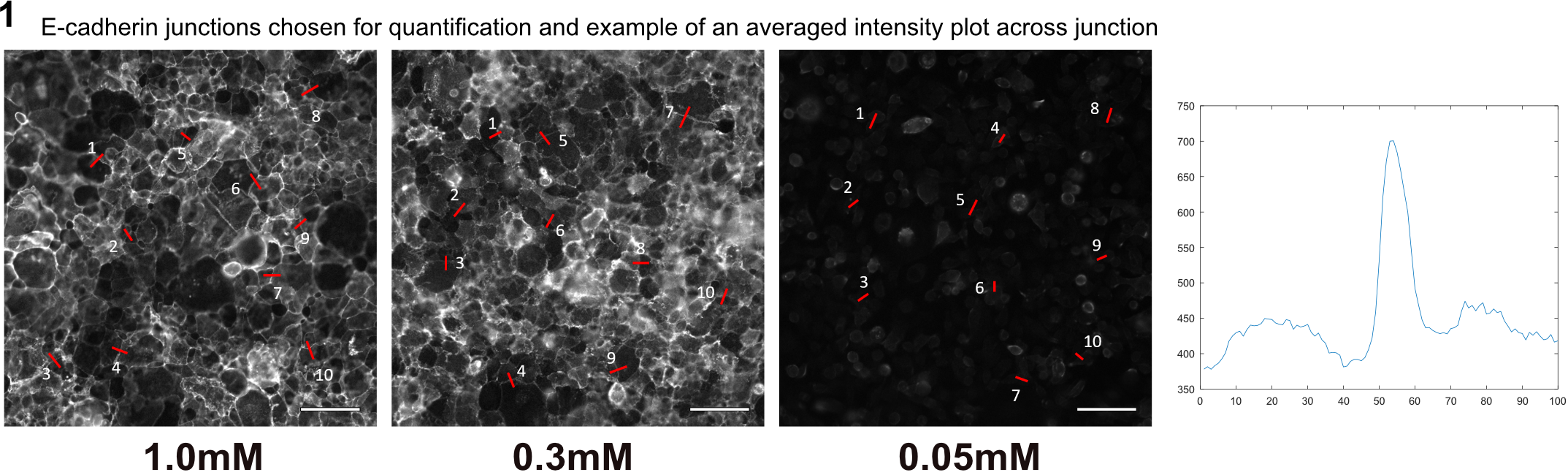

Fig. S1. Immunofluorescence imaging for E-cadherin with DECMA-1 for monolayers cultured in high (1.0mM), medium (0.3mM), and low (0.05mM) calcium media. Red lines indicate the cross-junctional lines across which the E-cadherin fluorescence intensity was quantified.

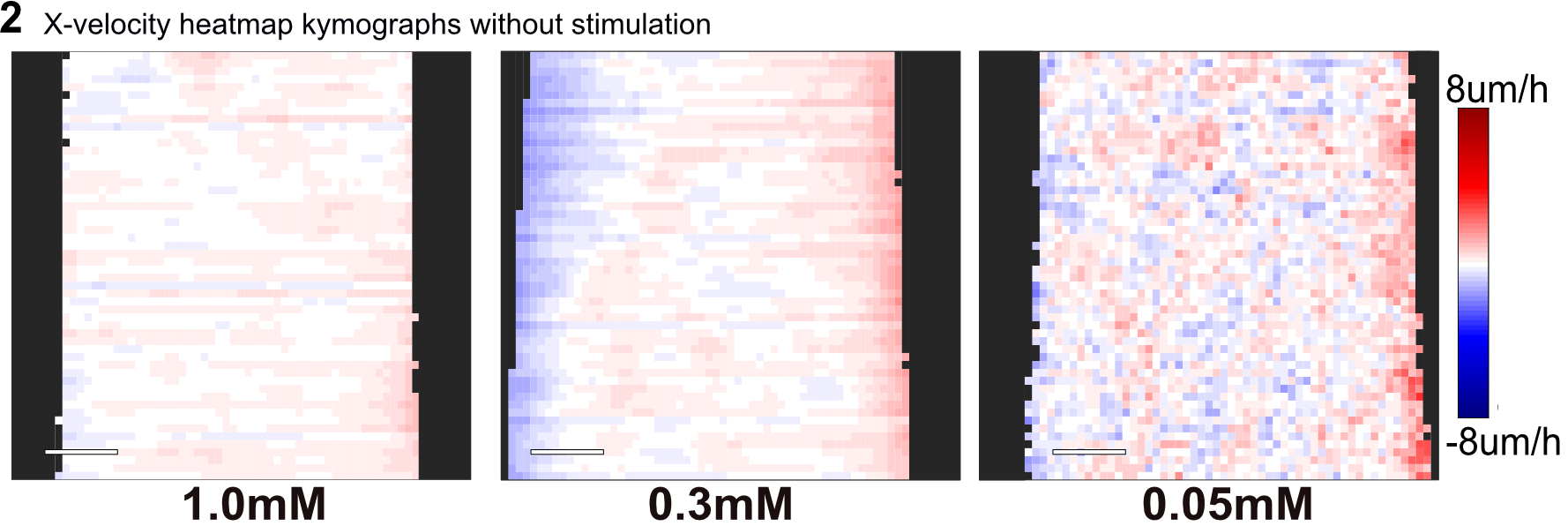

Fig. S2. X-velocity heatmap kymograph for varying calcium level without stimulation.

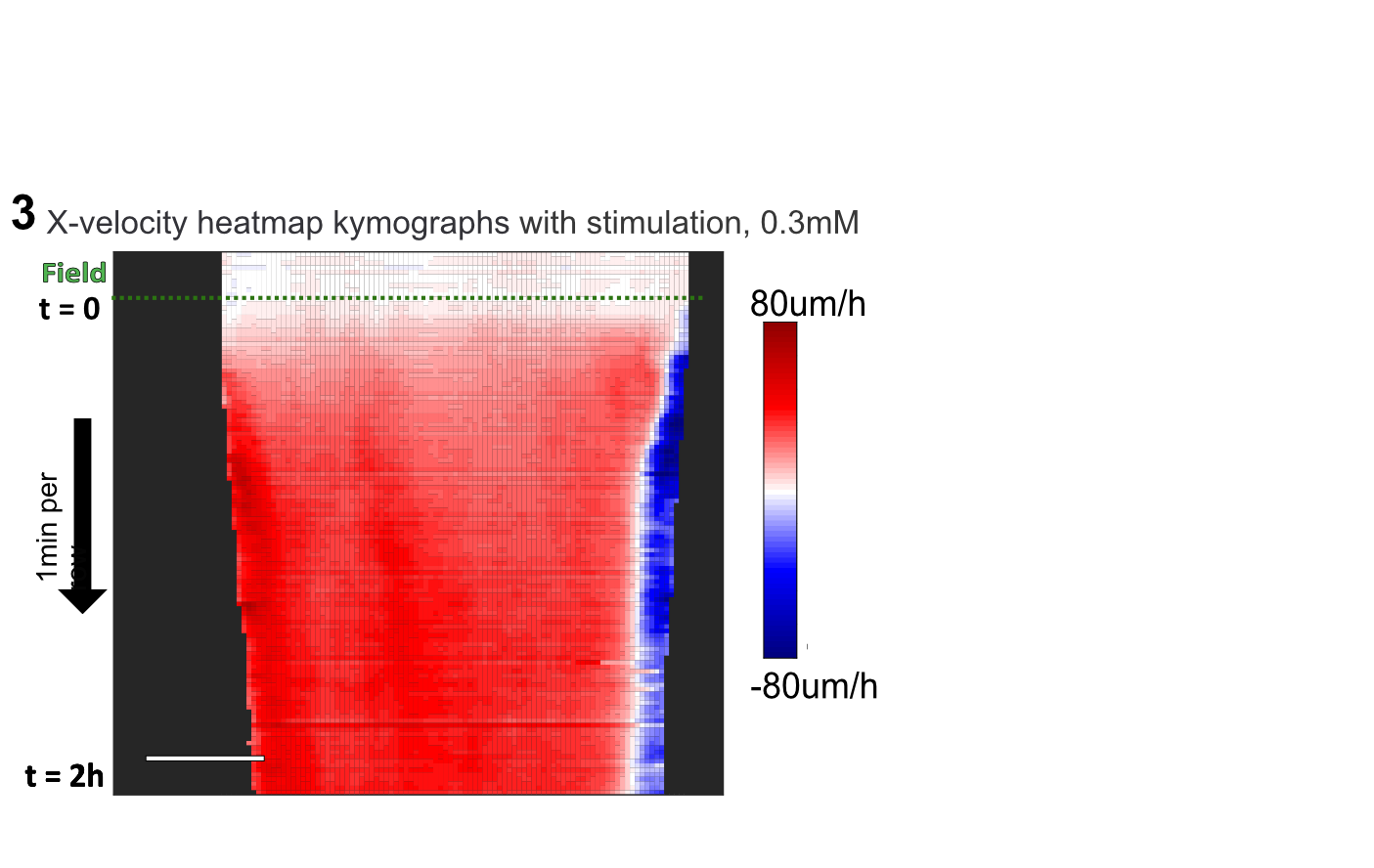

Fig. S3. X-velocity heatmap kymograph of electrotaxing keratinocyte monolayers cultured for 14 h in 0.3mM [Ca2+] media, stimulated at 2V/cm. Stimulation starts at dashed line. 1 min/ row. Scale bar = 500um.

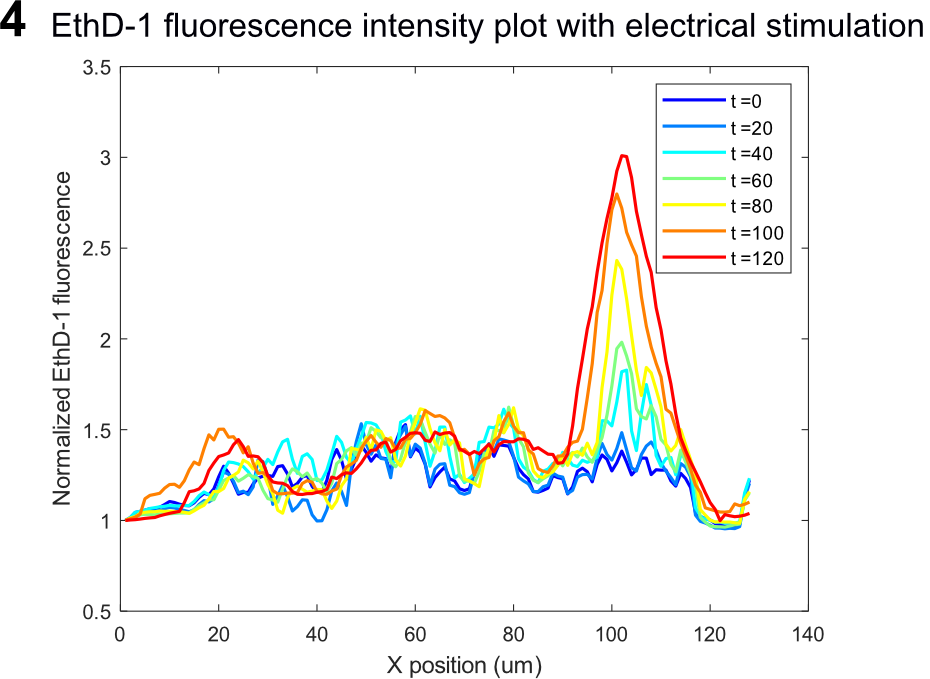

Fig. S4. Live-dead dye (EthD-1) fluorescence intensity across monolayer over 2h for keratinocyte monolayer cultured in medium calcium and stimulated at 2V/cm (N = 5). Monolayer migrates left to right, so the right edge is the leading edge.

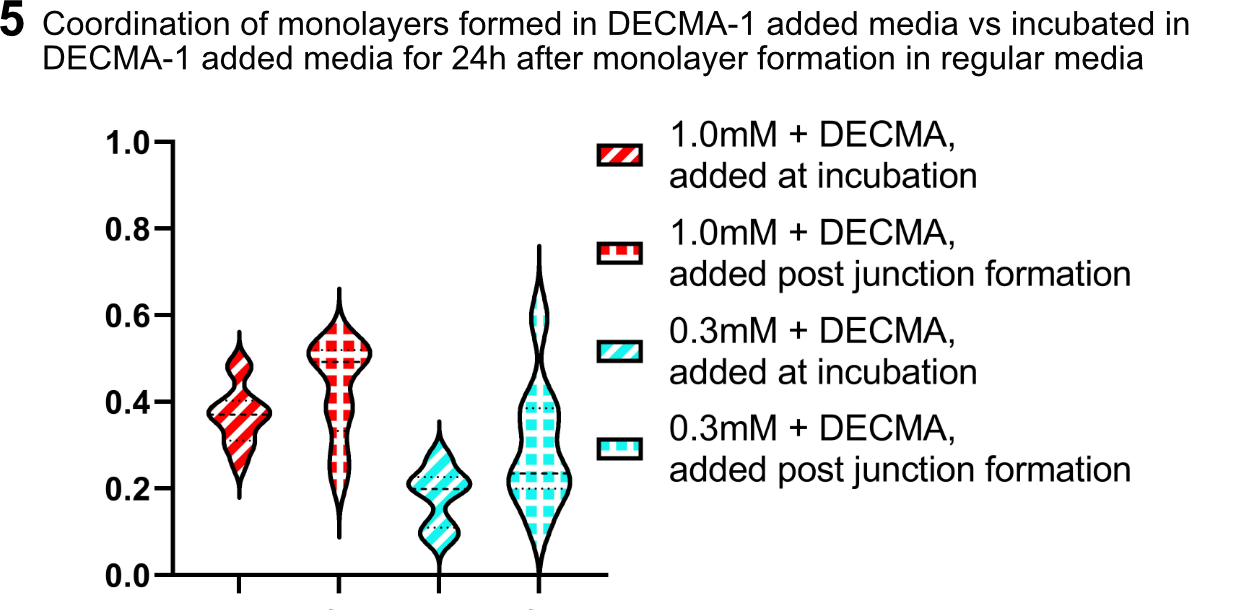

Fig. S5. Coordination values for monolayers formed with 50ug/mL DECMA-1 added at the start of the 14h incubation and monolayers treated with 50ug/mL DECMA-1 after incubation, post junction formation.

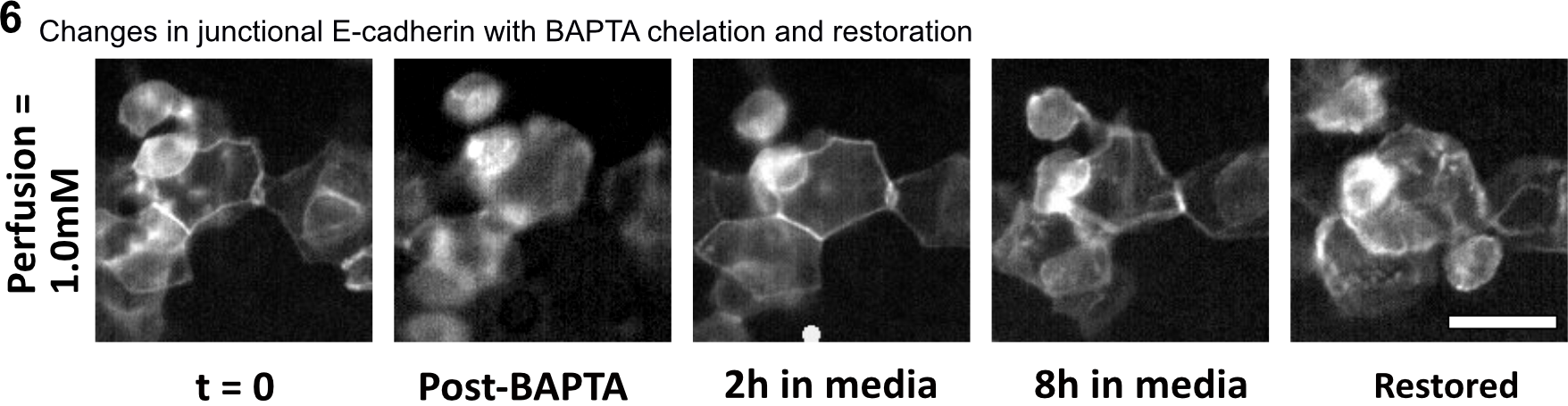

Fig. S6. E-cadherin fluorescence image of E-cadherin-GFP cells with treated with BAPTA, transferred to high calcium media, then incubated overnight in high calcium media. Top row: Image at t = 0, 1h BAPTA-treatment, 2h in high calcium media, 8h in high calcium media, and 14 h incubation in high calcium media. Scale bar = 20um.

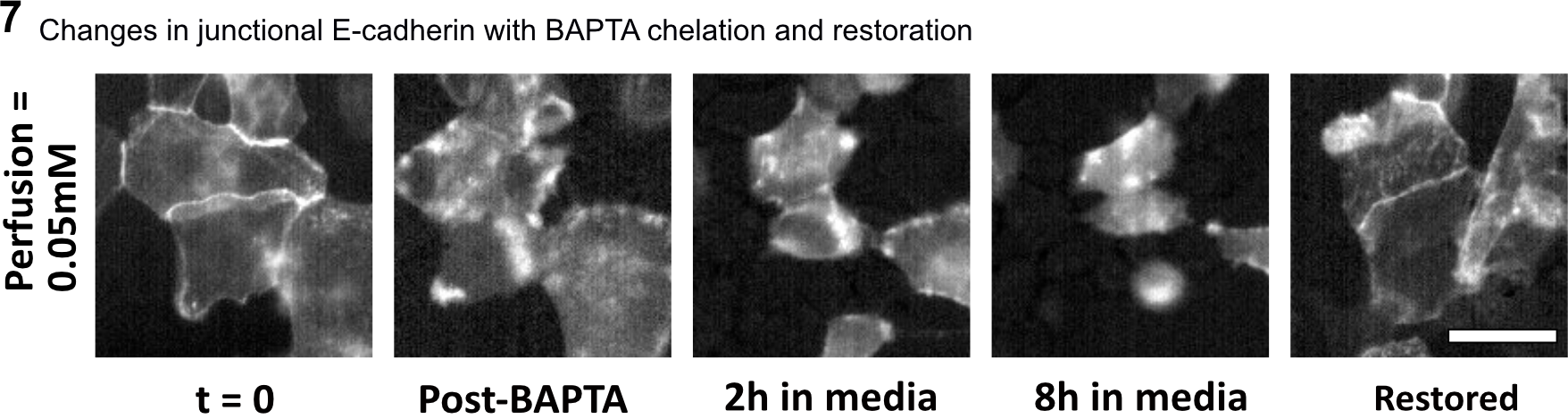

Fig. S7. E-cadherin fluorescence image of E-cadherin-GFP cells with treated with BAPTA, transferred to high calcium media, then incubated overnight in high calcium media. Images at t = 0, 1h BAPTA-treatment, 2h in low calcium media, 8h in low calcium media, and 14 h incubation in high calcium media. Scale bar = 20um.

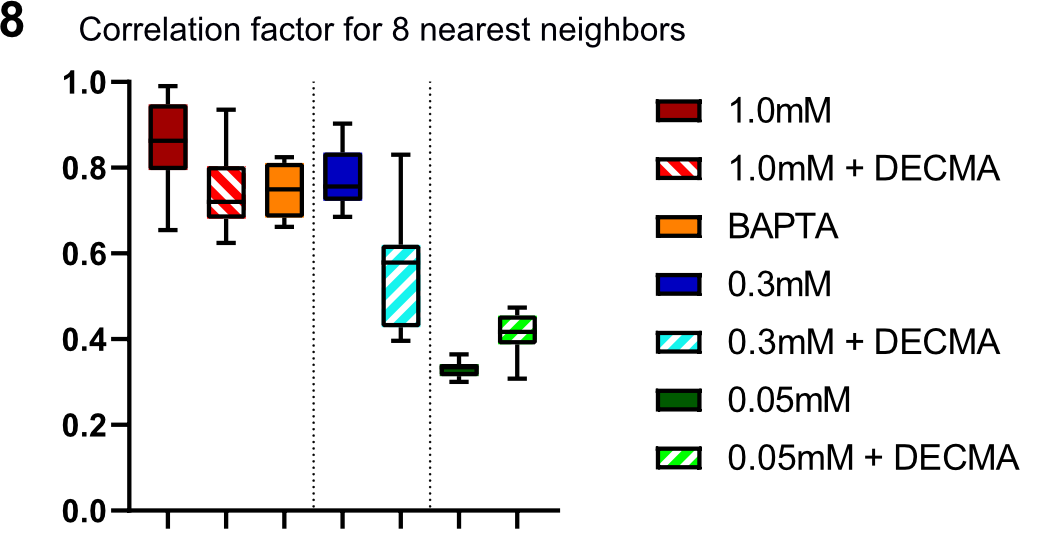

Fig. S8. Velocity correlation function values for 8 nearest neighbors (Methods) calculated for each experimental conditions.

**Supplementary movie legends**

Movie S1. Monolayers cultured and stimulated in extracellular medium of [Ca2+] = 0.05mM, 0.3mM, 1.0mM, no field. 10 min/frame. Scale bar = 500um.

Movie S2. Electrotaxis of monolayers cultured and stimulated in high (1.0mM), medium (0.3mM), and low (0.05mM) calcium media throughout (1h control, 8h stimulation) at 2V/cm. Electrical stimulation indicated with red circle. 10 min/frame. Scale bar = 500um.

Movie S3. Leading edge death using Live-Dead assay with EthD-1. Stimulation at 2V/cm with monolayer cultured and stimulated in medium calcium. 1 min/frame. Scale bar = 500um.

Movie S4. Single cells cultured and stimulated in high and medium calcium media. 10 min/frame. Scale bar = 100um.

Movie S5. Electrotaxis of monolayers cultured and stimulated medium calcium media, with 20uM blebbistatin and Y-27632 treatment. (1h control, 8h stimulation) at 2V/cm. Electrical stimulation indicated with red circle. 10 min/frame. Scale bar = 500um.

Movie S6. Electrotaxis of monolayers cultured and stimulated in high (1.0mM), medium (0.3mM), and low (0.05mM) calcium media with DECMA-1 treatment (1h control, 8h stimulation) at 2V/cm. Electrical stimulation indicated with red circle. 10 min/frame. Scale bar = 500um.

Movie S7. Electrotaxis of monolayers cultured high calcium medium, treated in 20uM BAPTA for 1h, and stimulated in high and low calcium media at 2V/cm. Electrical stimulation indicated with red circle. 10 min/frame. Scale bar = 500um.

Movie S8. Electrotaxis of monolayers cultured in high calcium media, treated with 20uM BAPTA for 1h, stimulated in low calcium media for 8h at 2V/cm, and restored without stimulation in high calcium media for 14h. Electrical stimulation indicated with red circle. 10 min/frame. Scale bar = 500um.

Movie S9. Convergent electrotaxis of monolayers cultured in high calcium media, treated with 20uM BAPTA for 1h, stimulated in low calcium media for 12h at 2V/cm, and restored without stimulation in high calcium media for 14h. Electrical stimulation indicated with red circle. 10 min/frame. Scale bar = 500um.
